# Effect of ozone exposure on Amyotrophic Lateral Sclerosis (ALS) pathology using a mice model of TDP-43 proteinopathy

**DOI:** 10.1101/2021.02.12.430915

**Authors:** Ana Rodriguez, Agueda Ferrer-Donato, Marta Cabrera-Pinto, Susana Seseña, Paloma Fernández, Alfonso Aranda, Carmen M. Fernandez-Martos

## Abstract

**Background:** Ozone (O_3_), one of the main photochemical pollutants in the atmosphere today, is a serious health risk factor. Although the effects of O_3_ exposure have been documented on many diseases, they have not yet been examined on Amyotrophic Lateral Sclerosis (ALS)- a fatal progressive and neurodegenerative disease.

**Objectives:** To investigate the effect of the O_3_ exposure in a mice model of TDP-43 proteinopathy, exploring a possible association between the O_3_ exposure and the ALS pathogenesis.

**Methods:** TDP-43^A315T^ and wild-type (WT) mice were exposed to O_3_ (0.25 ppm) or filtered air (FA) for 15 days (4 hours/day). We assessed (1) weight loss (2) motor performance (3) plasma glucose content and (4) metabolic markers from plasma samples of the animals.

**Results:** Throughout the experiment, we observed a progressive decline in body weight and the motor coordination in TDP-43^A315T^ mice compared to WT controls. Although there was a trend, there were no significant differences in the decline of body weight of TDP-43^A315T^ mice when exposed to either FA or O_3_. In O_3_-TDP-43^A315T^ mice, the disease duration lasted longer. In addition, O_3_-TDP-43^A315T^ mice showed improvements in motor performance as well TDP-43^A315T^ mice were hypoglycemic compared to WT mice. However, FA-TDP-43^A315T^ mice showed lower plasma glucose levels at the disease end-stage. We found altered levels of adipokines and metabolic proteins in TDP-43^A315T^ mice compared to WT controls. A positive correlation was found among GIP and glucagon compared to insulin concentrations in control mice. Interestingly, resistin, Gastric Inhibitory Peptide (GIP), Glucagon Like Peptide 1 (GIP-1) and insulin levels were higher in O_3_-TDP-43^A315T^ mice.

**Discussion:** We provide new evidence about a mechanistic link between O_3_ exposure and the improvement of the metabolic disturbances present in TDP-43^A315T^ mice. Further studies are needed to corroborate the obtained results as they warrant to understanding the underlying mechanisms.

## Introduction

Environmental pollution is considered an international public health issue with multiple facets (Manisalidis et al. 2020). As urbanization and industrialization have reached unprecedented proportions worldwide in our era, ambient air pollution has become one of the biggest public health hazards worldwide, accounting for about 4 million deaths per year (Hoffmann et al. 2020; Sacks et al. 2020). Among the most pertinent air pollutants, tropospheric ozone (O_3_) stands out for its harmful health and environmental effects, contributing to climate change as a greenhouse effect gas. O_3_ is a highly oxidative gas that is formed by the action of solar radiation from photochemical reactions of other pollutants (i.e. nitrogen oxides, and volatile organic compounds) emitted by vehicles and manufacturing (Humans 2016). The toxic effects induced by O_3_ are registered in urban areas all over the world, causing biochemical, morphologic, functional, and immunological disorders (Lippmann 1989). Epidemiological studies have also shown associations between O_3_ exposure and increased morbidity and mortality (Vicedo-Cabrera et al. 2020). In fact, at present, O_3_ is already causing 55.000 premature deaths due to long-term exposure annually in Europe (Orru 2019). Furthermore, if we take into account the fact that O_3_ formation is temperature-dependent, and considering the impact of the changing climate, we can expect up to an 11% increase in ozone-associated mortality in some countries in Central and Southern Europe in 2050 (Orru 2019).

Extensive investigation about O_3_-related pathophysiological sequelae clearly show that exposure to this pollutant causes and exacerbates respiratory (D’Amato et al. 2019; Manisalidis et al. 2020; Sokolowska et al. 2019) and cardiovascular diseases (Bourdrel et al. 2017; Manisalidis et al. 2020; Song et al. 2020). What is more, O_3_ exposure could also increase the incidence of chronic metabolic disorders, obesity and diabetes type II (Shore 2019; Thomson et al. 2018), which has become one of the greatest global health threats of the 21st century. Furthermore, in humans, this pollutant increases the release of stress hormones, cortisol or rodent corticosterone (Thomson et al. 2018), and alters the metabolic and endocrine response to the glucose challenge, which could also contribute to metabolic dysregulation (Thomson et al. 2018). Remarkably, a growing number of studies strongly suggests that certain neurological pathologies, such as Alzheimer’s disease (Bello-Medina et al. 2019a; Rivas-Arancibia et al. 2010) and Parkinson’s disease (Kremens et al. 2014; Ritz et al. 2016), might be associated with the O_3_ exposure through a variety of molecular and cellular pathways that either damage brain tissue directly, or lead to a predisposition to these neurological alterations (Genc et al. 2012). Epidemiological evidence supporting this association is still limited, and the underlying mechanisms are poorly understood. As suggested by different studies (Filippini et al. 2020; Povedano et al. 2018; Yu et al. 2014), some environmental and occupational risk factors, including air pollution, could be associated with the occurrence of Amyotrophic Lateral Sclerosis (ALS). However, even though evidence exists of a potential link between air pollution and ALS, to date, no study has investigated this association.

ALS is a motor neuron disease characterized by the selective and progressive loss of upper and lower motor neurons of the cerebral cortex, brainstem, and spinal cord (Tapia 2014). The result of this loss is a rapidly progressive paralysis, which ultimately leads to death within three to five years of symptom onset. The estimated prevalence of this fatal disease is 5 per 100,000 in the United States and approximately 2-3 people per 100,000 of the general population in Europe. ALS, therefore, represents one of the most challenging socio-economic problems of our future. The majority of patients suffer from sporadic ALS (sALS; more than 90%), in which multiple risk factors from gene-environment interactions contribute to the disease pathogenesis. In contrast, only a small subset of patients suffers from familial ALS (fALS; less than 10%) due to their associated genetic dominant inheritance factor (Zarei et al. 2015). No gene-conferring susceptibility to a certain environmental exposure has been established in ALS. Indeed, the recognized mechanisms by which mutations or gene-environment interactions cause the progressive degeneration and death of motor neurons have not been elicited yet. To date, only two epidemiology studies have investigated the possible association between O_3_ exposure and the risk of ALS (Myung et al. 2019; Povedano et al. 2018). Only smoking has been firmly established to increase the risk for ALS (Oskarsson et al. 2015).

It has already been demonstrated that O_3_ exposure increases the risk of acute respiratory distress syndrome (Rhee et al. 2019), and recently it has been found that it could induce chronic state oxidative stress (Bello-Medina et al. 2019b). Progressive neuromuscular respiratory failure is the most common cause of mortality in ALS patients (Chio et al. 2009), and, while it is unclear whether oxidative stress is a primary or a secondary cause of neurodegeneration in ALS, data from both human tissue and studies in animal models of ALS suggest that oxidative stress is a major contributory factor leading to chronic motor neuron death (Barber et al. 2006; Petri et al. 2012). Therefore, there is a great need for studies to identify O_3_ as a risk factor to this disease since a better understanding of environmental risk factors could help to reduce exposures and it is hoped to markedly reduce ALS incidence over time.

In this context, our research evaluates *in vivo* the effect of the O_3_ exposure in the well-validated TDP-43^A315T^ murine model of TDP-43 proteinopathy, which recapitulates several aspects of the human ALS, providing, to our knowledge, the first insights into the association between the exposure to the O_3_ and the pathogenesis of ALS. Our studies highlight the importance of tailoring the use of animal models to the experimental question being addressed, to ensure translatability of findings.

## Methods

### Animals

Both mutant TAR DNA-binding protein 43 (TDP43^A315T^) mice (Wegorzewska et al. 2009) and age and gender-matched non-transgenic littermates (WT controls) (Strain No. 010700, Bar Harbor, ME, USA) were used in this study. This mouse model of ALS was generated using the mouse prion promoter (Prt) and a cDNA encoding human TARDBP with an A315T mutation (hTDP-43^A315T^) associated with fALS and containing an N-terminal FLAG-tag (Wegorzewska et al. 2009). These mice accumulate cytoplasmic TDP-43 in the brain and spinal cord, including TDP-43 aggregates (Ke et al. 2015). They do not, however, develop neuronal loss or paralysis (Wegorzewska et al. 2009). For all the experiments, TDP-43^A315T^ mice were used as hemizygotes. The Tg progeny were genotyped at 21 days of age by detecting hTDP-43 transgene using standard tail genomic DNA PCR analysis according to the distributor’s protocol. To avoid the ambiguity associated with reported gender-related differences in mean survival time of TDP-43^A315T^ mice (Hatzipetros et al. 2014; Wegorzewska et al. 2009), we used only male mice. The animals were group-housed under standard housing conditions with a 12h light-dark cycle, and food and water *ad libitum*. TDP-43^A315T^ mice were fed jellified food in pellet form (D5K52 food. Rettenmaier Ibérica, Spain) to mitigate the intestinal dysmotility phenotype and sudden death (Herdewyn et al. 2014), and their general health was regularly checked. All experimental procedures were approved by the Animal Ethics Committee of the National Hospital for Paraplegics (HNP) (Approval No 26/OH 2018), in accordance with the Spanish Guidelines for the Care and Use of Animals for Scientific Purposes.

### O_3_ exposure

Forty-two-day-old TDP-43^A315T^ mice and WT controls were divided in two groups, and exposed to O_3_ (9/ group, n = 3 TDP-43^A315T^ mice vs. n = 6 WT mice) or filtered air (FA) (9/ group, n = 3 TDP-43^A315T^ mice vs. n = 6 WT mice). In all cases, animals were exposed 15 consecutive days (4 hours/ day) following the protocol described by Bello-Medina *et al.* (Bello-Medina et al. 2019a). O_3_ was generated from pure O_2_ with a BTM 802N generator and distributed in a Plexiglas chamber (50 × 35 ×35 cm) together with zero air at a total flow of 15 L/ min. O_3_ concentration of the ambient air in the chamber was kept constant at 0.25 ppm and was continuously monitored by an Environment O342M analyzer (Envea, France). FA was obtained by filtering regular air with activated charcoal to reduce O_3_ concentration to a minimum of (< 0.02 ppm). During the exposures, animals had free access to food and water, and their general health was regularly checked.

### Monitoring and behavioral assessments

To monitor disease progression and onset-stage (defined as the last day of individual peak body weight before gradual loss occurs) determination, body weight lost was measured and motor performance was evaluated using rotarod test. All mice were weighed and assessed three times per week until the disease onset-stage. After that mice were then checked daily in the morning until the disease end-stage (defined as the weight below 80% of the initial weight on each of three consecutive days).

The rotarod motor test was performed on all mice once a week (Dang et al. 2014), starting from the 6 weeks of age until the day of euthanasia. Animals were previously trained for three consecutive days and three times a day to promote the learning of the task. The accelerated protocol was applied for this motor monitoring as described previously by Mandillo *et al.* (Mandillo et al. 2008). In brief, mice were placed on a rotarod apparatus (Model 7650, Ugo Basile) at a speed of 4 rpm with acceleration up to 40 rpm over 300 s. Three tests were performed for each mouse with a minimal interval of 20 mins, and the average of the longest two performances was taken as the final result for analysis.

### Measurement of plasma glucose content

At disease end-stage (~ 95-100 ± 2 days), animals were terminally anesthetized with sodium pentobarbitone (140 mg/ kg) and the whole blood was collected by cardiac puncture (Bewick et al. 2009). Immediately, glucose levels were measured using a glucometer (FreeStyle Optium™ Neo H glucometer, Abbott Diabetes Care Inc, USA), previously calibrated for plasma glucose levels. The samples remaining after the glucometer measurements were immediately centrifuged at 3380 *g* for 10 min at room temperature to separate plasma samples (Togashi et al. 2016), which were immediately frozen on dry ice and stored at – 80 °C for later analysis.

### Measurement of metabolic markers plasma

Adipokines (ghrelin, resistin and leptin) and metabolic biomarkers of insulin resistance from plasma samples were both analyzed by duplicate using the Bio-PlexPro mouse Diabetes group from Bio-Rad (Ref. 171F7001M), in a Luminex^®^ 200™ technology as previously described (Ortega Moreno et al. 2020). Samples were processed following the manufacturer’s instructions. The final concentration value of each metabolic marker was the result of the mean from the two duplicated measures.

### Statistical analysis

All data are presented as means ± standard error of the mean (SEM). Differences between means were assessed by two-way ANOVA followed by Tukey post hoc analysis. When only two groups were analyzed, a two-tailed Student *t*-test was used (equal variances assumed). For multiplex assays, the median for each sample group was determined for all the analytes, and Kruskal–Wallis test was performed. Spearman correlation was performed for both the adipokines and metabolic proteins, respectively. For all statistical tests, a *p* value of < 0.05 (CI 95%) was assumed to be significant. Statistical analysis was performed using GraphPad Prism software (version 8.3.1).

## Results

### Evaluation of O_3_ exposure on disease progression

To evaluate the effect of O_3_ exposure on disease progression, we monitored body weight loss and motor performance until the disease end-stage in WT controls and TDP-43^A315T^ mice exposed to O_3_ or FA for 15 days beginning at 42 days of age (on week 6). As TDP-43^A315T^ mice exhibit weight loss during disease progression (Esmaeili et al. 2013; Guo et al. 2012; Hatzipetros et al. 2014; Medina et al. 2014), we assessed the capacity of O_3_ exposure to modify weight changes over time. For this reason, the mice were weighed three times per week until the disease onset-stage. Then, the mice were weighed daily, and an average of the weights for each week was evaluated. Starting on week 8. During the 15 days of exposure, no differences in weight gain between groups (O_3_ group vs. FA group) were displayed (Figure 1). A two-way ANOVA with repeated measures revealed a significant genotype interaction (Figure 2A; *F*_(15, 81)_=87.72, *p*<0.0001), indicating a sustained decline in body weight in TDP-43^A315T^ mice compared to WT controls in responses to FA or O_3_ over time. However, although there was a trend, no significant differences were observed between FA-exposed or O_3_-exposed TDP-43^A315T^ mice, indicating that O_3_ exposure had no significant effect on weight loss in TDP-43^A315T^ mice. In addition, the calculation of the disease onset using this parameter indicated that TDP-43^A315T^ O_3_-exposed develop symptoms later than FA-treated TDP-43^A315T^ mice. Using body weight measurements (criterion of 10% body weight loss) an average onset of 75 ± 5 days of age was determined for in TDP-43^A315T^ mice in responses to FA, whereas in responses to O_3_, TDP-43^A315T^ mice presented a phenotype at 83 ± 1 days of age (Figure 2B). To further assess changes in weight due to the effect of O_3_ exposure on disease progression based on the determined disease onset, we calculated disease duration (Figure 2C). Average disease duration of the animal was calculated as the time between the onset of disease and the day of death. Interestingly, while there were no significant differences in the decline of body weight in TDP-43^A315T^ mice in responses to FA or O_3_ over time, comparatively the disease duration was longer in TDP-43^A315T^ mice in responses to O_3_ (Figure 2C), although O_3_-TDP-43^A315T^ mice did not live significantly longer compared with FA-TDP-43^A315T^ mice.

**Fig. 1.**
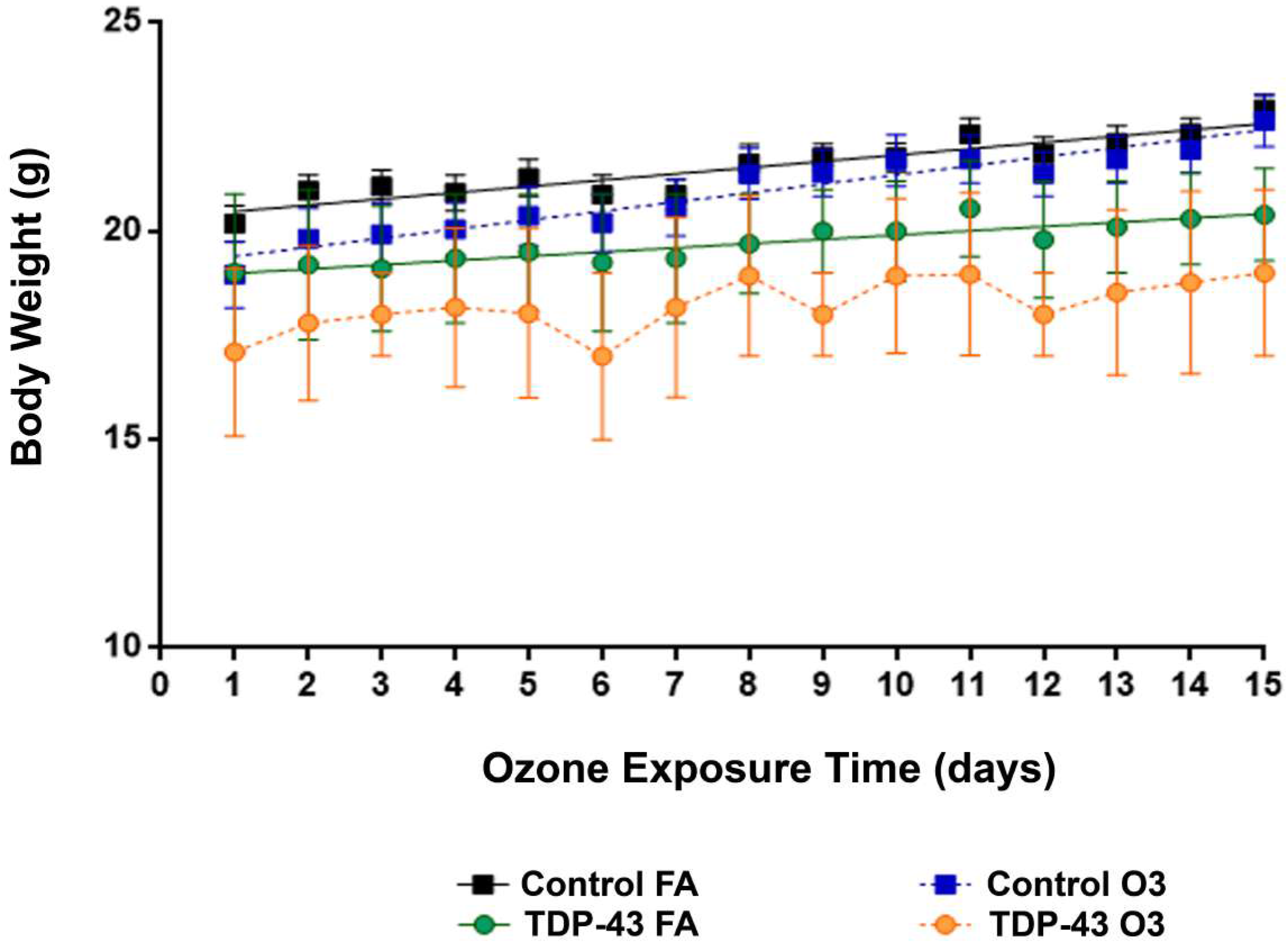
O_3_ exposure beginning at the asymptomatic state of disease does not alter body mass in TDP-43^A315T^ mice. Body weight was monitored daily in WT controls and TDP-43^A315T^ mice exposed to FA or O_3_ for 15 days, beginning at the asymptomatic state, at 42 days of age (~ 6 weeks of age). No differences in weight gain between groups was determined. Values are expressed as a mean ± SEM. Comparison between groups was performed by two-way ANOVA followed by Tukey test, unless stated otherwise. Abbreviations: Control; non-transgenic littermates (WT controls), TDP-43; TDP43^A315T^ mice; FA, filtered air, O_3_; ozone. Corresponding graph as control-FA (n =6, black square and solid line), control-O_3_ (n = 6, blue square and dashed line), TDP-43^A315T^-FA (n = 3, green circles and solid line), TDP-43^A315T^-O_3_ (n = 3, orange circles and dashed line).

**Fig. 2.**
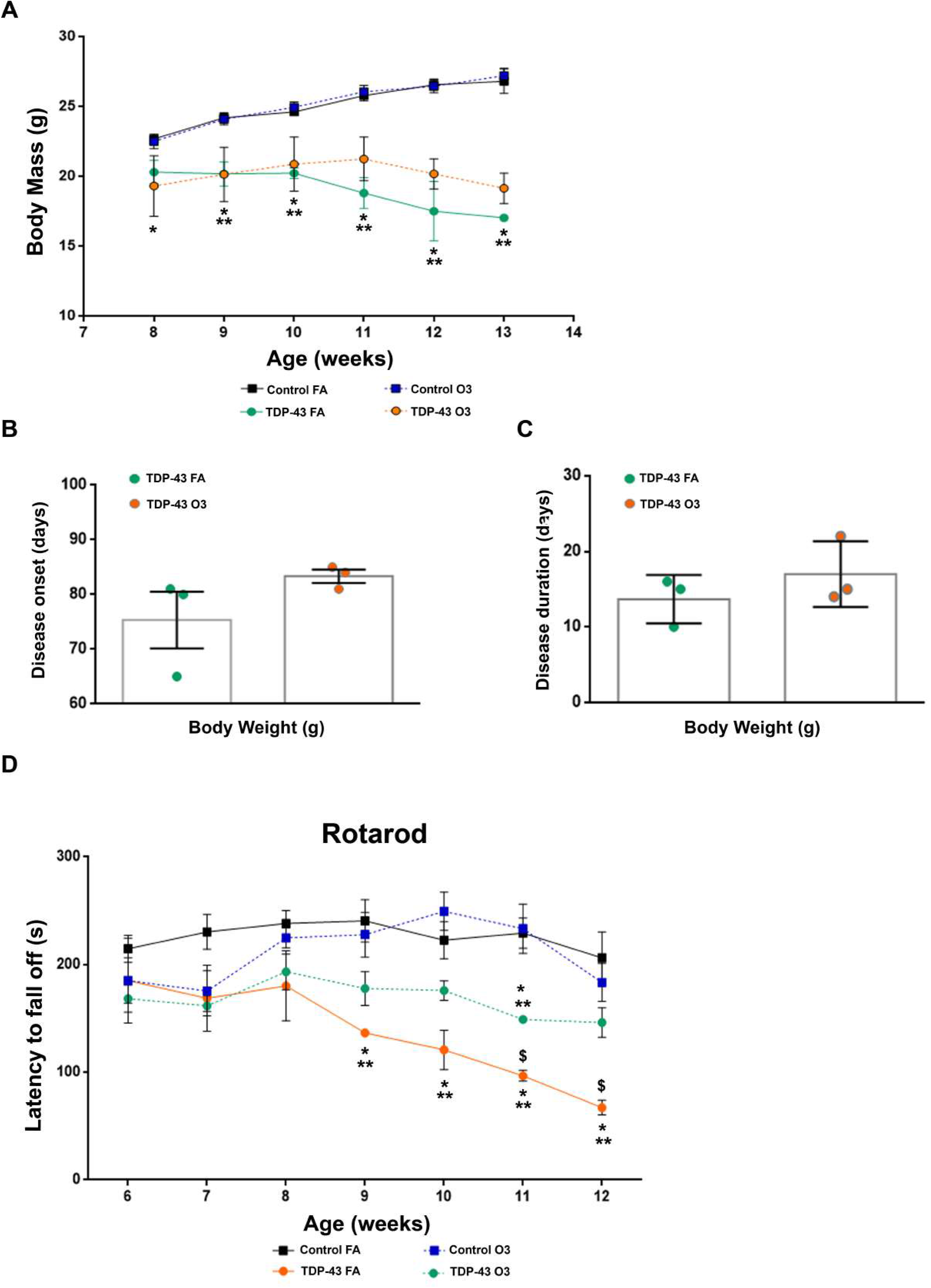
O3 exposure beginning at the asymptomatic state of disease does not alter body mass change but significantly improves motor performance in TDP-43^A315T^ mice. **(A)** Body weight was monitored over time in WT controls and TDP-43^A315T^ mice exposed to FA or O_3_. Starting weight on week 8. No significant differences were observed between FA-exposed or O_3_-exposed TDP-43^A315T^ mice. **(B)** Average disease onset and disease duration **(C)** was determined in WT controls and TDP-43^A315T^ mice exposed to FA or O_3_ using body weight as a physiological parameter. Comparatively the disease duration was higher in TDP-43^A315T^ mice in responses to O_3_. **(D)** Behavioral assessment of motor function was performed in WT controls and TDP-43^A315T^ mice exposed to FA or O_3_ over time. Significant differences between FA- and O_3_-exposed mice were seen. Values are expressed as a mean ± SEM. Comparison between groups was performed by two-way ANOVA, where ^*^ *p <* 0.05 vs. FA-exposed WT control mice; ^**^*p <* 0.05 vs. O_3_-exposed WT control mice; ^&^ *p <* 0.05 vs. O_3_-exposed TDP-43^A315T^ mice. Abbreviations: Control; non-transgenic littermates (WT controls), TDP-43; TDP43^A315T^ mice; FA, filtered air, O_3_; ozone. Corresponding graphs as per A., i.e. control-FA (n = 6, black square and solid line), control-O_3_ (n = 6, blue square and dashed line), TDP-43^A315T^-FA (n = 3, green circles and solid line), TDP-43^A315T^-O_3_ (n = 3, orange circles and dashed line).

Furthermore, we also tested motor behavior to determine if O_3_ exposure beginning during the asymptomatic stage of the disease could alter disease phenotype. We analyzed the time spent on an accelerating rotarod starting from the age of 6 weeks. Lower scores on the rotarod are indicative of impaired locomotor function. Our results indicate a progressive decline in motor coordination in TDP-43^A315T^ mice, confirming the progressive motor deficits of the TDP-43^A315T^ mouse model reported previously (Stallings et al. 2010). A two-way ANOVA with repeated measures revealed a significant interaction of the group by week (Figure 2D; *F*_(6,97)_=3.051 *p*<0.008), indicating differential change over time in rotarod performance. However, O_3_-exposed TDP-43^A315T^ mice showed a significant improvement in motor performance at later time points, indicating the effect of O_3_ exposure on motor function. Indeed, the rotarod test displayed that FA-exposed TDP-43^A315T^ mice suffer a more significant drop in performance and progressive impairment in motor capacity over time.

Overall, although O_3_ exposure had no significant effect on disease-associated weight loss in TDP-43^A315T^ mice, our results indicate that TDP-43^A315T^ mice exposed to O_3_ developed a significant motor and coordination improvement.

### Evaluation of O_3_ exposure in plasma glucose levels

We next aimed to determine if plasma glucose levels, were altered in TDP-43^A315T^ mice exposed to O_3_ or FA as imbalanced energy homeostasis is a consistent finding in different ALS mice (Kim et al. 2011). O_3_ exposure had a significant effect on plasma glucose levels of WT controls (221.5 ± 8.596 mg/ dL vs. 167.2 mg/ dL ± 14.26; *p<0.004*) (Figure 3). In addition, our results indicated that with either FA or O_3_-exposed, TDP-43^A315T^ mice were hypoglycemic compared to WT mice at the disease end-stage (Figure 3), confirming the disturbances in the energy metabolism of the TDP-43^A315T^ mouse model reported previously (Chiang et al. 2010). Interestingly, FA-TDP-43^A315T^ mice showed lower plasma glucose levels at later time points (96 mg/ dL ± 13.39 vs. 142.3 mg/ dL ± 30.14, *p<0.18*) (Figure 3), indicating that TDP-43^A315T^ mice exposed to O_3_ required less energy demand in the late symptomatic stage.

**Fig. 3.**
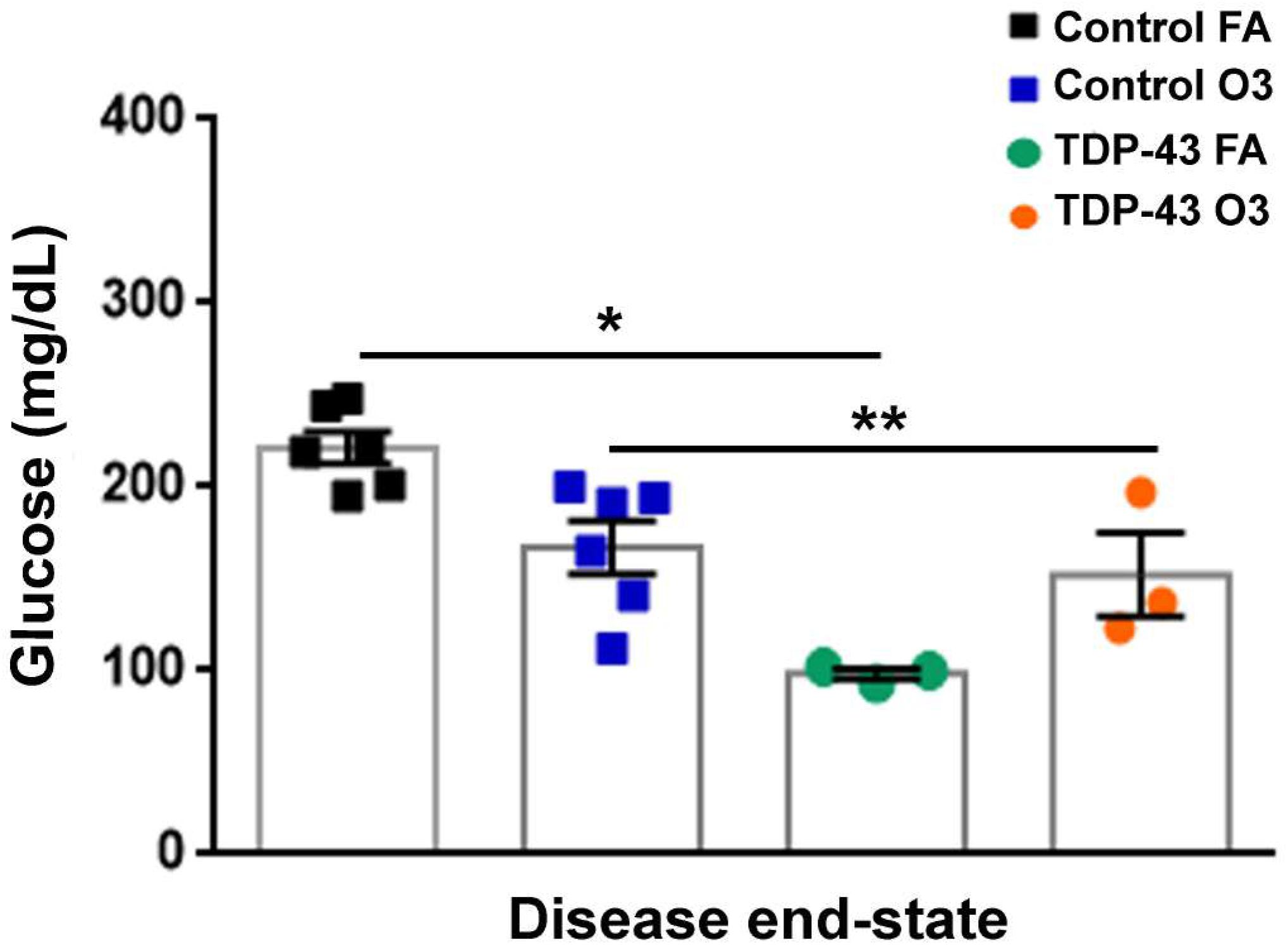
O_3_ exposure beginning at the asymptomatic state increased plasma glucose levels in TDP-43^A315T^ mice. Plasma glucose concentration was measured in WT controls and TDP-43^A315T^ mice exposed to FA or O_3_ at disease end-stage (~ 95-100 ± 2 days). Quantification revealed the highest levels in response to O_3_ exposure in TDP-43^A315T^ mice. In addition, significant differences between FA- and O_3_-exposed WT mice were seen. Values are expressed as a mean ± SEM. Comparison between groups was performed by two-way ANOVA, where \**p <* 0.05 vs. FA-exposed WT control mice; \*\**p <* 0.05 vs. O_3_-exposed WT control mice. Abbreviations: Control; non-transgenic littermates (WT controls), TDP-43; TDP43^A315T^ mice; FA, filtered air, O_3_; ozone.

### Evaluation of O_3_ exposure in adipokines and metabolic proteins

To evaluate the effect of O_3_ exposure on the adipose tissue, serves as the body’s major energy storage compartment, and secretes numerous endocrine mediators, we measured the levels of adipokines that influence metabolism. The expression of ghrelin, resistin and leptin in TDP-43^A315T^ mice and WT controls is presented in Table 1. We found altered levels of these adipokines in the plasma of TDP-43^A315T^ mice compared to WT controls. The concentration of resistin in plasma was significantly higher in O_3_-exposed TDP-43^A315T^ mice, however, there was no difference in levels of ghrelin and leptin between TDP-43^A315T^ mice exposed to either O_3_ or FA (Table 1). In addition, O_3_ exposure had a significant effect on the plasma concentration of leptin in WT mice. However, no correlation was found among the serum adipokines levels in WT controls or TDP-43^A315T^ mice exposed to FA or O_3_, respectively (Figure 4A, B). Moreover, for metabolic proteins, we observed altered levels of GIP, GLP-1, glucagon, and insulin in the plasma of TDP-43^A315T^ mice relative to WT controls (Table 2). Comparatively, plasma levels of GIP, GLP-1, and insulin are higher in O_3_-exposed TDP-43^A315T^ mice relative to FA-exposed TDP-43^A315T^ mice (Table 2), though no statistical differences were determined. It is worth noting that a positive correlation was found among the plasmatic levels of GIP and glucagon compared to insulin concentrations in WT controls (Figure 5A, B).

**Table 1.** Adipokines concentration in the groups of the study and Kruskal–Wallis comparison (n = 16). Values were reported as median (interquartile range); Kruskal–Wallis comparisons were performed. In the case of leptin, Dunn’s post hoc test was significant only for Control-O_3_ vs. TDP-43-FA (*p* = 0.012). Abbreviations: Control; non-transgenic littermates (WT controls), TDP-43; TDP43^A315T^ mice; FA, filtered air, O_3_; ozone.

| Molecule (pg/ml) | Control FA | Control O<sub>3</sub> | TDP-43 FA | TDP-43 O<sub>3</sub> | *p* |
| --- | --- | --- | --- | --- | --- |
| Ghrelin | 6.25 | 10.75 | 7.60 | 7.00 | 0.56 |
| Resistin | 11.50 | 11.75 | 3.60 | 8.33 | 0.01 |
| Leptin | 10.75 | 14.25 | 4.40 | 4.66 | 0.0001 |

**Table 2.** Metabolic biomarkers of insulin resistance concentration in the groups of the study and Kruskal–Wallis comparison (n = 16). Values were reported as median (interquartile range); Kruskal–Wallis comparisons were performed. Abbreviations: Control; non-transgenic littermates (WT controls), TDP-43; TDP43^A315T^ mice; FA, filtered air, O_3_; ozone.

| Molecule (pg/ml) | Control FA | Control O<sub>3</sub> | TDP-43 FA | TDP-43 O<sub>3</sub> | *p* |
| --- | --- | --- | --- | --- | --- |
| GIP | 5.62 | 7.87 | 8.00 | 14.00 | 0.12 |
| GLP-1 | 6.37 | 6.12 | 10.20 | 11.67 | 0.30 |
| Glucagon | 5.25 | 7.62 | 11.40 | 9.16 | 0.28 |
| Insulin | 5.62 | 8.00 | 9.50 | 11.33 | 0.45 |
| PAI-1 | 5.00 | 7.50 | 10.40 | 11.33 | 0.25 |

**Fig. 4.**
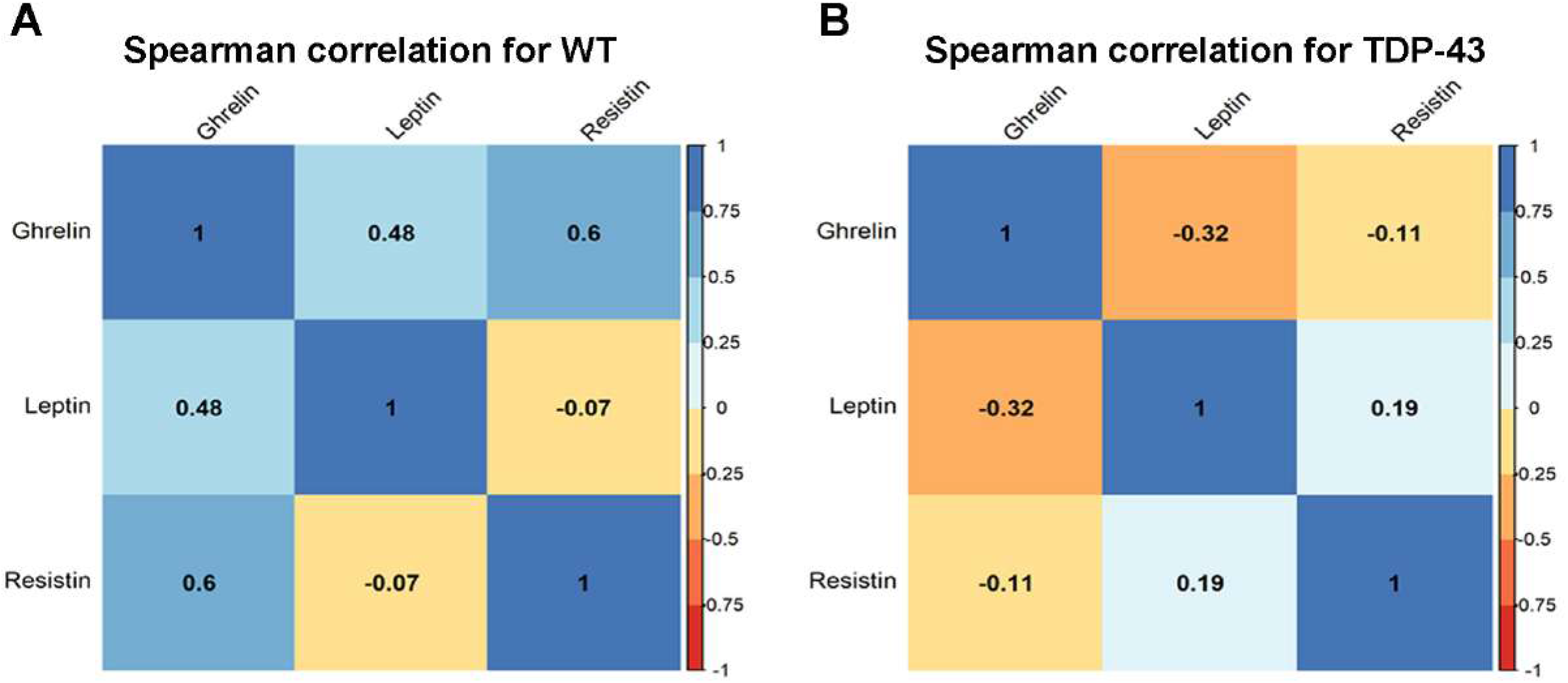
Spearman correlations for the adipocytokines ghrelin, resistin and leptin in WT controls and TDP-43^A315T^ mice exposed to FA or O_3_. In this figure, the red and blue squares refer to negative and positive correlations, respectively; \**p*<0.05. The color intensity of the squares is proportional to the correlation coefficient. In the legend at the right size, the intensity of the color shows the rate of correlations and the corresponding relationships.

**Fig. 5.**
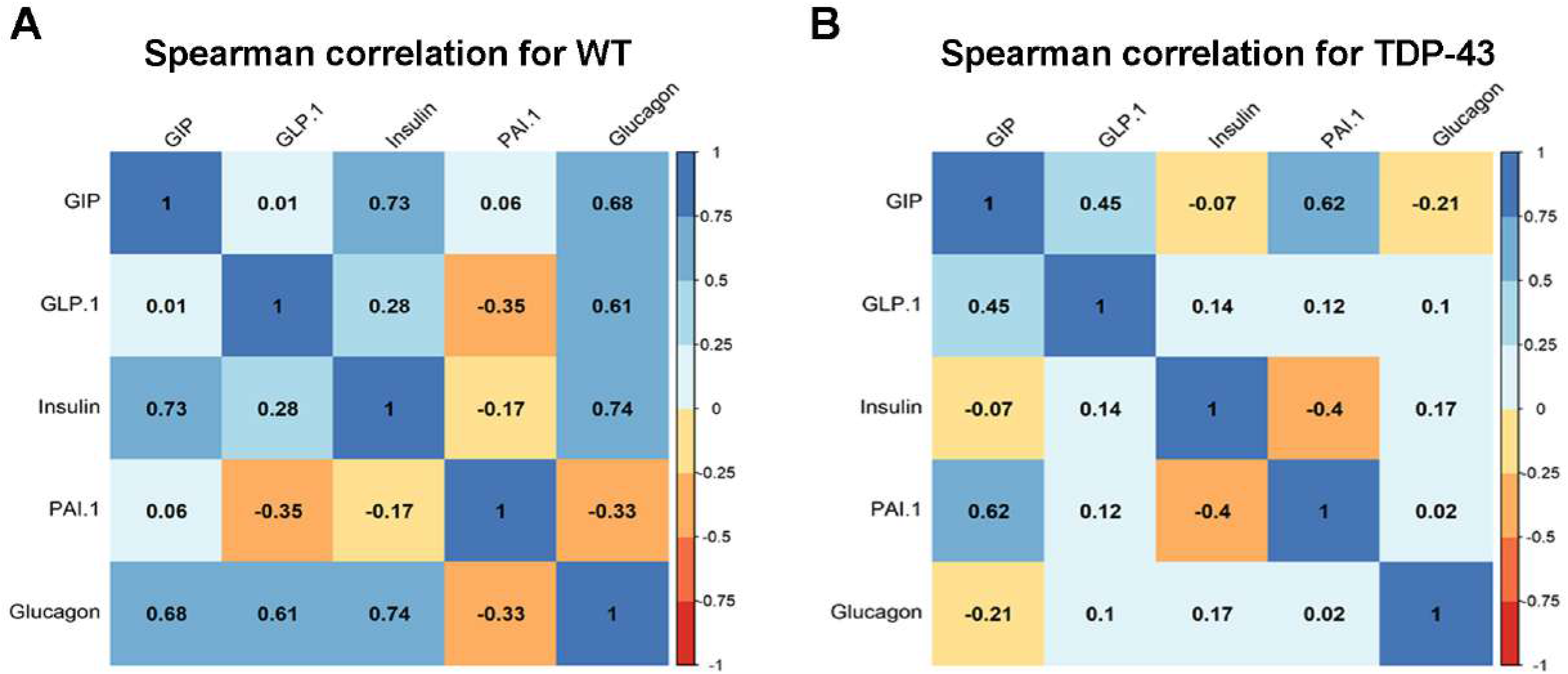
Spearman correlations for metabolic proteins in WT controls and TDP-43^A315T^ mice exposed to FA or O_3_. In this figure, the red and blue squares refer to negative and positive correlations, respectively; \**p*<0.05. The color intensity of the squares is proportional to the correlation coefficient. In the legend at the right size, the intensity of the color shows the rate of correlations and the corresponding relationships.

## Discussion

Despite recent, significant progress in the ALS research field, no cure has yet been discovered to stop the neurodegenerative progression of this disease or to improve the lives of ALS patients. The underlying pathogenesis is not fully understood, and therefore, the disease can be very difficult to identify in its earlier stages resulting in poor and late diagnosis. It is believed that an interaction between genetic and non-genetic factors, including air pollutants such as O_3_, may be involved in the development of ALS. However, epidemiological evidence supporting this association is very limited and poorly defined, and whether O_3_ exposure can modify underlying disease-related pathological changes is unclear. In this study, we sought to investigate the effects of O_3_ exposure in TDP-43^A315T^ mice, which develop neuropathology and behavioral deficits similar to human ALS. Unexpectedly, our results displayed that O_3_ exposure did not alter disease onset, progression, or survival of TDP-43^A315T^ mice. In fact, we found that O_3_ exposure significantly improves motor and coordination performance of TDP-43^A315T^ mice and resulted in beneficial changes to plasma glucose levels at disease end-stage. Surprisingly despite previous studies suggesting the association between exposure to air pollutants and increased susceptibility to ALS (Filippini et al. 2020; Povedano et al. 2018; Yu et al. 2014), overall, our study supported the potential benefit of O_3_ in modifying ALS-like pathology and disease phenotypes in mice.

Nevertheless, extensive investigations about O_3_-related pathophysiological sequelae clearly show that exposure to this pollutant exacerbates respiratory diseases, such as asthma, respiratory infections, and chronic obstructive pulmonary disease (D’Amato et al. 2019; Manisalidis et al. 2020; Sokolowska et al. 2019). The adverse effect of O_3_ on cardiovascular diseases has been also well established (Bourdrel et al. 2017; Manisalidis et al. 2020; Song et al. 2020). However, the impact of this pollutant on metabolic alterations is less understood, even when the metabolic-impairment effects of O_3_ have been determined in humans (Shore 2019). In this context, our results demonstrated that O_3_ exposure had a significant effect on the regulation of the expression of certain adipokines and metabolic proteins that are linked to metabolic disease (i.e. obesity and diabetes type II) in WT control mice. This is concordant with previous mouse studies that saw that acute O_3_ exposure causes endocrine and metabolic changes, increasing food intake and body fat mass (Nappi et al. 2016). Interestingly, we have also identified specific alterations in levels of metabolic proteins such as GIP, PAI-1, and insulin in response to O_3_ exposure in TDP-43^A315T^ mice. This is also important to consider in the case of O_3_ exposure since it has been shown that PAI-1 and insulin are metabolic proteins associated with an increase in adiposity and body mass index (BMI) (Kahn et al. 2006; Ngo et al. 2015). Remarkably, GIP stimulates insulin secretion in response to food intake (Elliott et al. 1993). These findings suggest that O_3_ exposure might cause endocrine and metabolic changes in TDP-43^A315T^ mice, though the mechanism is currently unknown, which would help reduce disturbances in energy metabolism associated with the progression of ALS. Accordingly, our study indicated that O_3_ exposure mitigates the sustained decline in body weight in TDP-43^A315T^ mice over time. Certainly, this result is very interesting considering patients with ALS are unable to maintain their body weight, in part because of the decline in their nutritional conditions (Bouteloup et al. 2009; Desport et al. 2001). Indeed, rapid weight loss in patients with ALS disease is associated with worse disease outcomes (Ahmed et al. 2019). Likewise, disturbances in energy metabolism have been associated with the progression of human ALS (Vandoorne et al. 2018). Therefore, it would be plausible that this effect of O_3_ exposure on body weight gain in TDP-43^A315T^ mice could be due to increased food intake. However, there was no difference in levels of ghrelin and leptin, two appetite-stimulating hormones (Klok et al. 2007), between FA or O_3_-exposed, TDP-43^A315T^ mice. In contrast to these results, we find that O_3_ exposure had a significant effect on the plasma concentration of leptin in WT controls. Likewise, a positive correlation was found among the plasmatic levels of GIP and glucagon relative to insulin concentrations in this group, supporting previous experiments conducted in rodents, which strongly associated exposure to O_3_ with the development of obesity and diabetes type II (Campolim et al. 2020).

It may also be worthwhile to consider other potential disease-relevant effects of O_3_ exposure, for example, in the TDP-43^A315T^ mice, in which a progressive motor impairment has been described (Stallings et al. 2010). In our study, we observed that O_3_ exposure improved the progressive decline in motor coordination in TDP-43^A315T^ mice over time, and resulted in higher plasma glucose levels at later time points, raising the hope that reducing energy demand could offer benefits in ALS disease. The majority of ALS patients have glucose metabolism defects (i.e. impairments in glucose transport, glycolysis, among others) (Tefera et al. 2021), which are likely to contribute to disease. Indeed, impaired glucose homeostasis in ALS could occur, at least in part, because of higher levels of circulating glucagon (Hubbard et al. 1992). Our data indicated that TDP-43^A315T^ mice exposed to O_3_ developed motor and coordination improvement, correlating with a delay on the disease duration, and the premature death of characteristic of TDP-43 mice expressing hTDP-43^A315T^ transgene (Esmaeili et al. 2013; Guo et al. 2012; Hatzipetros et al. 2014; Medina et al. 2014). In addition, we found lower levels of circulating glucagon in O_3_-exposed TDP-43^A315T^ mice compared to FA-treated TDP-43^A315T^ mice, and a positive correlation was determined among the plasmatic levels of GIP and glucagon compared to insulin concentrations in WT controls. Remarkably, O_3_ exposure had a significant effect on plasma glucose levels of WT controls, suggesting that the possible causative role that O_3_ plays in the development of insulin resistance was not attributed to excessive energy demand alone. Therefore, since it has been suggested an increase in disease severity in ALS patients is correlated with increased circulating glucagon levels (Ngo et al. 2015), our data are congruent with the observations that insulin resistance is related to disease severity and outcome in ALS (Muddapu et al. 2020).

In summary, our findings show that O_3_ exposure could offer benefits to ALS patients. To date, there are only two medications, riluzole (glutamate antagonist) (Miller et al. 2002) and edaravone (free radical scavenger) (Rothstein 2017), approved by the US Food and Drug Administration (FDA) Unfortunately, none of these treatments reverse the damage already caused by ALS. Indeed, no cure is available for ALS to stop the neurodegenerative progression of ALS or to improve the lives of patients suffering from this terrible disease. Hence, it is imperative to find an effective therapy to minimize the sufferings of these patients. Our study provides the first preliminary evidence of a mechanistic link between O_3_ exposure and the improvement of the metabolic disturbances present in TDP-43^A315T^ mice. How O_3_ exposure contributes to disease progression of TDP-43-related disease remains unclear. Further studies are required to investigate whether O_3_-exposed TDP-43^A315T^ mice show alterations in levels of adipokines and metabolic proteins with an important role in regulating food intake and energy balance in the brain at late stages of the disease.

## Acknowledgments

The authors would like to gratefully acknowledge the Animal Facility and Experimental Surgery Unit of the UDI-HNP for their excellent technical support. Also, we extend our gratitude to Dr Nicolas Valiente-Parra, Department of Biosciences, University of Oslo (Norway) for his assistance with the statistical analysis. This work was supported by the funding from the Consejería de Educación, Cultura y Deportes, Fondo Europeo de Desarrollo Regional (FEDER), Junta de Comunidades de Castilla-La Mancha (SBPLY/17/180501/000303).

